# HLA-SPREAD: A Natural Language Processing based resource for curating HLA association from PubMed abstracts

**DOI:** 10.1101/2021.01.05.425409

**Authors:** Dhwani Dholakia, Ankit Kalra, Bishnu Raman Misir, Uma Kanga, Mitali Mukerji

## Abstract

Extreme complexity in the Human Leukocyte Antigens (HLA) system and its nomenclature makes it difficult to interpret and integrate relevant information for HLA associations with diseases, Adverse Drug Reactions (ADR) and Transplantation. PubMed search displays ∼144,000 studies on HLA reported from multiple diseases in diverse locations. Currently, IPD-IMGT/HLA database houses data on 28,320 HLA alleles. We developed an automated pipeline with a unified graphical user interface HLA-SPREAD that provides a structured information on SNPs, Populations, REsources, ADRs and Diseases information. Information on HLA was extracted from ∼24 million PubMed abstracts extracted using Natural Language Processing (NLP). Python scripts were used to mine and curate information on diseases, filter false positives and categorize to 24 tree hierarchical groups and named Entity Recognition (NER) algorithms followed by semantic analysis to infer HLA association(s). This resource from 112 countries and 32 ethnic groups provides interesting insights on: markers associated with allelic/haplotypic association in autoimmune, cancer, viral and skin diseases, transplantation outcome and ADRs for hypersensitivity. Summary information on clinically relevant biomarkers related to HLA disease associations with mapped susceptible/risk alleles are readily retrievable from HLASPREAD. The resource is available at URL http://hla-spread.igib.res.in/. This resource is first of its kind that can help uncover novel patterns in HLA gene-disease associations.

## INTRODUCTION

Human Leukocyte Antigen (HLA) locus consists of six classical genes (HLA-A, -B, -C, -DP, -DQ and -DR) that play an important role in eliciting immune response against pathogens (1) and three non-classical genes (HLA-E, -F and -G) that interact with Natural Killer cells to regulate virus-infected and malignant cells (2). HLA genes harbour a large number of mutations. As of September 2020, there are 28,320 HLA alleles reported in IPD-IMGT/HLA database. These variations mostly arise to generate defensive mechanisms against pathogens. However, some variations also confer risk to autoimmune diseases like rheumatoid arthritis, multiple sclerosis, Type 1 diabetes and Graves’ disease etc. More than 100 different autoimmune diseases, infectious diseases and adverse drug reactions have been reported to be associated with HLA genes (3–5). These alleles have clinical utility as diagnostic markers for example in rheumatoid arthritis, ankylosing spondylitis (6–8). They are also used in genetic screening e.g. HLA-B*57:01 in Caucasian population for abacavir hypersensitivity, HLA-B*15:02 in Chinese and Asians for carbamazepine induced life-threatening conditions like Stevens-Johnson syndrome (SJS) and toxic epidermal necrolysis (TEN) (9, 10). In the context of transplantation, mismatch of HLA alleles between donor and recipient impacts the solid organ and hematopoietic stem cell transplantation outcomes (11, 12). Each of the reported studies is unique in itself as they describe the molecular basis of disease associations, HLA matching and anti-HLA antibody formation that are relevant for transplantation. Besides, studies also report some relevant and associated clinical information, e.g different HLA-B27 subtypes are reported to be associated with clinical categories under spondyloarthropathies (13). There are other studies that implicate HLA allele association with the composition of gut microbiome and diseases (14–16). The expanse of this information is immense as there is wide genetic variability and heterogeneity among populations (17). Although advancements in HLA typing technologies has been beneficial in identifying novel HLA sequences (18), this has also led to reporting the same HLA allelic variant using different HLA nomenclature.

With the rapid increase in biomedical data, HLA alleles and their associations in multiple diseases, it becomes imperative to create a platform with structured information to query and retrieve relevant information. Current knowledge about HLA limits to individual papers that can be searched through PubMed or reviews where a subset of studies has been summarised. Hitherto, there exists no database that complies the existing HLA related information in an organised framework. In absence of such a repository, resource sharing among researchers and clinicians becomes a big challenge.

The integration of computer sciences with biomedical research has accelerated the progress, both in terms of novel discoveries and data structuring. Natural Language Processing (NLP) is a method to extract relevant information from unstructured data (19). A simple NLP pipeline contains 4 components: data assembly, pre-processing and normalization, Named Entity Recognition (NER) and Relation Extraction (RE). The output of NLP algorithms, i.e. structured dataset can be used to generate insights via direct interpretation or through downstream analyses. In recent times, NLP methods have started gaining popularity in biological sciences. For instance, Rakhi et.al (20) reported a text mining pipeline to study spice-disease associations and link phytochemicals from different spices/herbs to diseases. Another report by Lee et.al highlights BioBERT, a pre-trained biomedical language representation model that can be used for various text mining tasks like NER, RE and question answering, specifically on biomedical datasets. Similarly, PubTator Central (21) is an open access tool available via NCBI that uses text mining algorithms for assisted bio-curation of entities in literature. The tool uses NER to identify and thus highlight six bio-entities viz. Gene, Disease, Chemical, Mutation, Cell Line and Species from abstracts and open access articles available on PubMed. Another interesting report by Kuleshov et.al(22) presents a machine compiled database for studying genotype-phenotype associations generated using applications of text mining on genome-wide association studies (GWAS). All these resources work on similar text mining algorithms, but each has a different set of applications and tasks to perform. The use of these resources as such in addressing the HLA research often overlooks the extent of variability of HLA complex and involved parameters in this domain. For instance, PubTator Central is able to mine gene names from literature, but would not pick HLA allele information e.g., it will highlight HLA-DRB1 when the user search query is HLA-DRB1*01:01. Conventional processes to individually mine a large amount of unstructured literature available on HLA research requires both manpower and resources. For understanding and integrating the observations from HLA studies we require knowledge of genomic datasets, i.e. diseases, SNPs, drugs, populations, and ethnic groups along with an understanding of the relationship between them. NLP based text mining is an ideal approach to understand the complexity of this process to create a structured information.

We provide HLA-SPREAD (**Figure 1**) as a platform for integrated HLA resources that has been developed using NLP to understand the complexity of this locus. The resource provides a platform to summarize HLA related genomics knowledge as well as to design and develop new hypothesis. In this study, we have used publicly available ∼24 million peer reviewed abstracts. We extracted biomedical entities including HLA alleles, diseases, SNPs, drugs and geographical locations. We also tried assigning positive and negative relationships between disease and alleles. This HLA connectivity was then used to address biologically and clinically relevant objectives like HLA-biomarkers and risk and protective alleles for various diseases.

**Figure 1.**
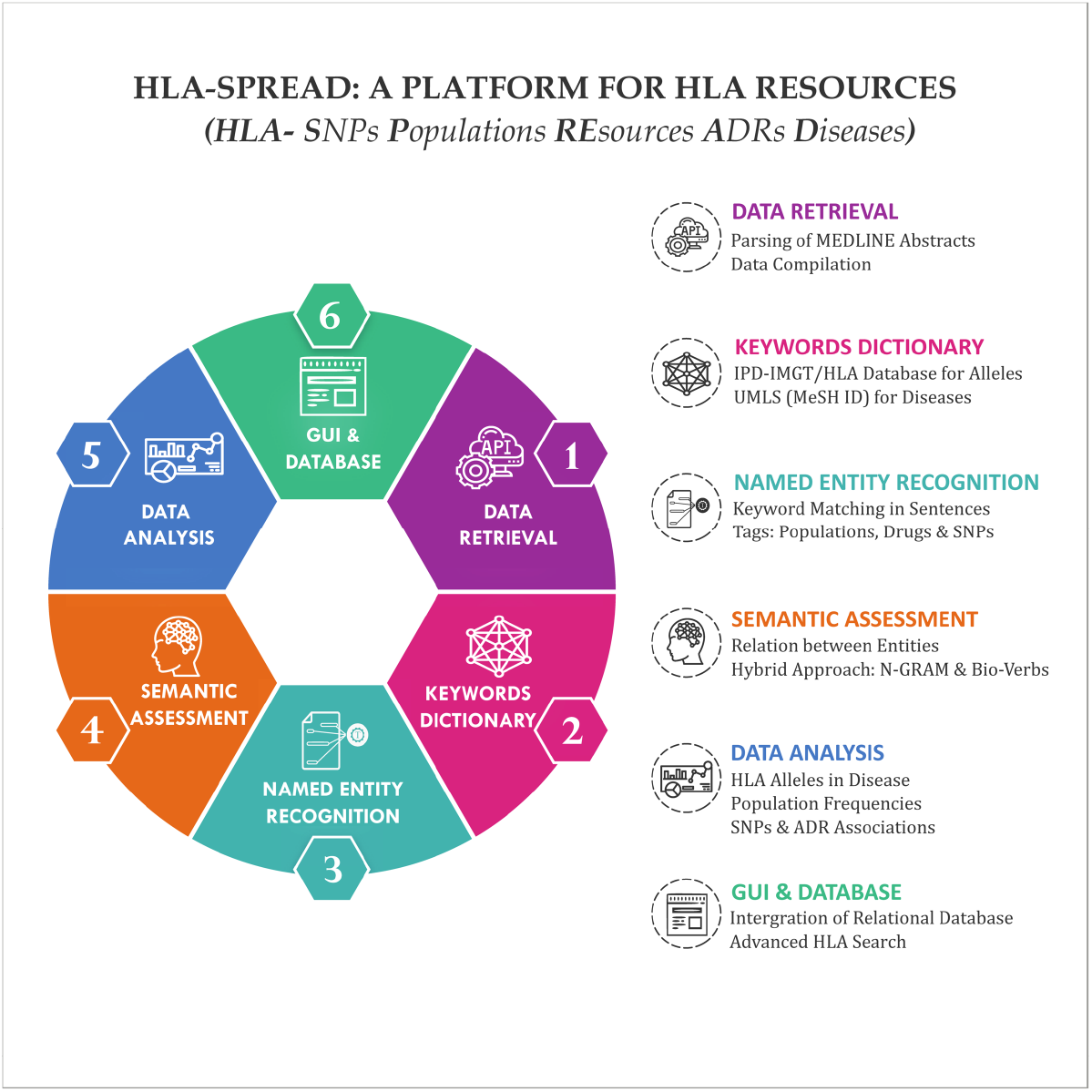
Workflow of HLA-SPREAD: An automated pipeline developed to extract information related from ∼110,000 studies related to HLA retrieved from over 24 million abstracts. Structured information from these abstracts was created using Natural Language Processing methods developed into a database HLA-SPREAD. The various resources used at each step are indicated.

## MATERIAL AND METHODS

### Data Retrieval

MEDLINE was used as a source of biomedical literature that comprises more than 24 million peer-reviewed articles from over 5600 scholar journals. Bulk data was downloaded from the FTP server in XML format. HLA alleles with nomenclature were downloaded from IPD-IMGT/HLA database(23). To maintain uniformity in disease names and their IDs, we used MeSH keywords from UMLS (Unified Medical Language System). Drugs associated with side effects were obtained from SIDER 4.1 and Allele Frequency Net Database (AFND) (24, 25). Allele frequency of HLA alleles were also taken from AFND. Extensive Pre-processing was done on all the datasets before they were implemented in the pipeline.

### Pre-processing and Keywords Dictionary

#### PubMed parsing

A modified version of PubMed parser was used to extract PMID, title, abstract, publication date, journal, article type and authors’ information from MEDLINE biomedical literature dataset (26). Only records with the above information were considered for further analysis and stored in a tabular format. All the subheadings in the abstract viz background, introduction, objective, method, experimental design, result, discussion, importance, setting, design, study objective, patients, participants and conclusion were removed.

#### Disease Dictionary

Mentions of disease keywords were identified using a dictionary created from UMLS 2019MRCONSO.RRF (27). UMLS is a set of biomedical vocabulary that includes data from OMIM, Gene Ontology, Clinical repositories, Medical Subject Headings (MeSH) and NCBI taxonomy. In this study, we used MeSH descriptors including Entry Term (ET), Main Heading (MH), Preferred Entry term (PEP), Descriptor Sort Version (DSV) and Machine Permutation (PM). Descriptor Entry Version (DEV) was excluded as keywords belonging to this category were incomplete, e.g. abdominal injury was reported as abdominal inj. These descriptors are assigned a unique MeSH ID which is stored in a hierarchical format with 24 head categories along with a unique Descriptor ID. We termed the root form of the disease as level-zero and top-level diseases as level-one for our analysis. Multiple forms of a disease like diabetes insipidus, diabetes mellitus, type 1 diabetes, juvenile-onset diabetes and others are assigned the same MeSH ID. This dataset was also supplemented with keyword variants such as plural and lemmatised forms to increase the search space.

#### HLA Dictionary

Keywords for HLA alleles and their nomenclature were fetched from the centralized repository of international ImMunoGeneTics project (IMGT) database. IMGT is updated quarterly with submission or deletion of alleles and their nomenclature and currently houses 28,320 alleles. Many reports do not follow the conventional HLA allele nomenclature which makes mapping a strenuous task. To maximally capture all HLA alleles, we created a dataset comprising of all possible keywords including the removal of special characters, whenever required. We have also attempted mapping all the old nomenclature to the current allele names. This dictionary also includes few generic HLA keywords like HLA class I, HLA class II, HLA linked and HLA associated. There are few alleles based on old nomenclature that belong to more than one antigenic group, hence they were put under “broad antigen” category. A few haplotypes that were a combination of more than one HLA allele were grouped in “haplotype” category.

### Named Entity Recognition

#### Keyword Matching across Abstracts

A python-based NER pipeline was implemented to filter abstracts based on a dictionary matching approach using parallel multiprocessing. Disease and HLA allele keyword dictionaries were used for initial screening. Abstracts were converted to lower case with special characters removed and if a match was found in either title or text, the abstract was sentence tokenized using sentence tokenizer, a part of python Natural Language Tool Kit (NLTK). We encountered a great extent of variability in the names of disease keywords. Most of it had special characters like (-) and (‘) in the keyword or with the plural and singular forms. To deal with the former, we kept instances of sentences where special characters were not removed, this increased the search space that enables capturing of keywords such as Stevens-Johnson syndrome (Stevens Johnson syndrome), Graves’ disease (Graves disease). Our disease dictionary was already enriched with plural and lemmatized forms of keywords to tackle the latter. For HLA allele keywords, word boundary-based regex matching was implemented to search alleles in the sentences. Sentences with at least a single mention of both HLA allele and disease keywords were considered for further steps.

### Identification of Tags: Populations, Drugs and SNPs

#### Populations

The filtered abstracts were processed using spaCy NLP tagging algorithm (model: en_core_web_md) to search for mention of populations in text. From the two output tags, i.e. GPE (Geo-Political Entities) and NORP (Nationalities Or Religious Groups), we selected the keywords having the latter as GPE tag often reported scientific names of organisms as populations when applied on biomedical data, e.g. scientific names such as Chlamydia spp. and Chlamydomonas spp. were reported under GPE tags. The output was classified into countries and ethnic groups for further analysis with the help of an expert anthropologist. Manual curation of the obtained list was also done to remove plural and inappropriate entries.

#### Drugs

The information on drugs with side effects were taken from the SIDER database (SIDER 4.1). We also added 16 drugs from AFND, whose information was missing in SIDER. The list of drugs was mapped across the dataset to check for its occurrences in selected HLA related abstracts. There were many instances where drug names were subpart of disease keywords, e.g. “insulin” was obtained as a false match wherever it was present as a part of the disease name “insulin dependent diabetes mellitus”. A small python snippet was written to remove such false positives.

#### SNPs

SNP IDs were mapped across abstracts of the HLA dataset using the RegEx module of python. The algorithm iteratively searched for all instances of RSIDs using regular expression “[rR][sS][0-9]{2,}”. All the tags captured in various sentences of abstracts were stored in a list of strings format along with their respective PMIDs for facilitated future access.

### Semantic Assessment

#### N-GRAM Evaluation and Manual Labelling

N-grams refers to a contiguous sequence of n items (can be syllables, letters, or word pairs) in a text for determining the context of said items in a sentence or paragraph. We used the functions of NLTK viz. WordNetLemmatizer, WordPunctTokenizer and CollocationFinder to create a corpus of NGRAMS (n=1, 2 and 3) from the abstract dataset. After removal of stop words, that do not add significant meaning to the context, a subset consisting of all reported verb/adverb(n=1), adverb-verb(n=2,3) combinations based on a frequency cut-off was filtered out using Part of Speech (POS) tags of tokenised words. We observed that N-grams for negative labels often gave misleading information, e.g. “HLA-B27 negative” refers to the absence of allele rather than a negative association between entities. Hence, we used very stringent criteria for choosing negative labels. Manual annotation of positive and negative labels was then carried out on this dataset and a total of 1127 labels (**Supplementary Table 1**) were categorised (1107 positive and 20 negative) for labelling the sentences. We assert a positive label where the HLA allele is positively associated with disease and hence its presence makes individuals susceptible to disease, whereas in negative statements the HLA allele is negatively associated with disease and hence protective for the disease. We also considered negation words like “not, none, no” which if present, can reverse the actual meaning of the sentences. Instances of above mentioned three keyword sets (positive, negative and negation) were iteratively searched in all the sentences. Further, a coding scheme was constructed using the binary layout to label sentences as positive, negative, negation, complex ambiguous. Sentences having no match from either of the categories were labelled as others.

#### Root-Verb and Associated Adverbs using Dependency Parsing

Dependency parsing refers to the formation of a tree layout based on the semantics of a sentence, where the root node is represented by a verb that describes relation between different entities of that sentence. Direct implementation of word tokenization, first step in dependency parsing, generates multiple tokens for single allele and disease keywords as shown in **Figure 2**. Therefore, to ensure the accuracy of the algorithm, the allele and disease keywords present in each sentence were replaced with @GENE and @DISEASE tags and a parse tree was then generated using StanfordCoreNLP python module (Stanford-corenlp-full-2018-10-05 package). The list of verbs obtained from the root nodes of all the sentences in the dataset was manually curated under positive and negative labels. We also added a category “Studied/Investigatory” that doesn’t convey any positive or negative context but have mentions of both entities together, e.g. “To investigate the association of HLA-A, B, and DRB1 alleles with leukaemia in the Han population in Hunan province”.

**Figure 2.**
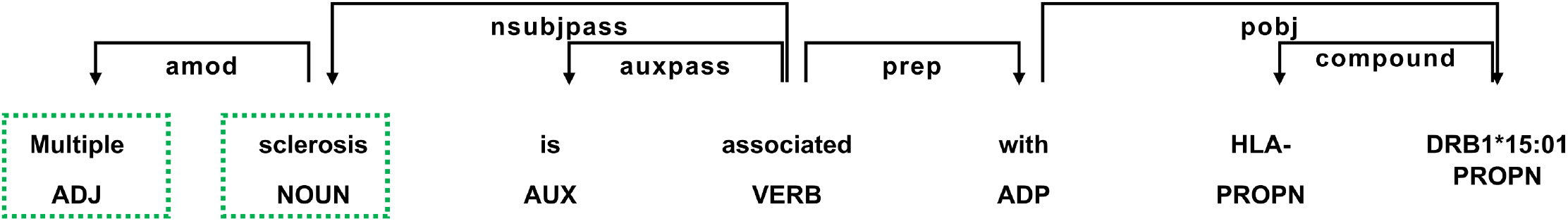
Tokenization and Dependency parsing: In this example, the keyword “Multiple sclerosis” is tokenized to ”Multiple” and “sclerosis” separately with parts of speech Adjective and Noun respectively.

#### Sentence Annotation

We termed our approach as “hybrid approach” for labelling sentences, where annotation was done using both N-gram labels and the type of root verbs. If a sentence had a positive N-gram label and a positive root verb, that inferred the relationship between entities as associated or linked, then the sentence was labelled as positive. For negative labelling also we used the same approach. Finally, labelling of sentences were grouped into different categories: 1) Positive, 2) Negative, 3) Both positive and negative, referring as Complex sentences, 4) Positive/negative + negation referring as Ambiguous group, 5) Investigatory and 6) Others (-)

#### Database and web server

HLA SPREAD database is built for quick and easy retrieval of information related to HLA genes. The web interface was designed in HTML5, CSS3 & ES6 (JavaScript) and the backend was developed in Laravel 8 (PHP Web Framework) & MySQL for the database. Laravel is a PHP web framework proposed for the development of a web system following the Model View Controller (MVC) architecture. We used D3.js for data visualization and SQL indexing for search table integration. The server was hosted using Apache HTTP (PHP) server. The database uses Relational Database Management System with data stored in the table. JavaScript handles the data visualizations and Laravel handles the search queries, indexing, and the data export section. This web interface is compatible with various devices and browsers except the feature “Show entries” in the search tab is visualised best in Mozilla Firefox.

## RESULTS

### Mining Medline literature for HLA association

NLP based text mining of 24 million publicly available biomedical abstracts provided 41845 abstracts with either one or more sentences that describe the relationship between the HLA alleles and diseases. To understand the distribution of various kinds of articles published among the filtered abstracts, we studied the article type per year trend from 1975 to 2019 (**Figure 3**). We found research journal, comparative study and review articles to have maximum numbers every year. In addition, there were papers corresponding to clinical trials phase I, II, III and IV and observational studies highlighting the importance of this locus in translational studies.

**Figure 3.**
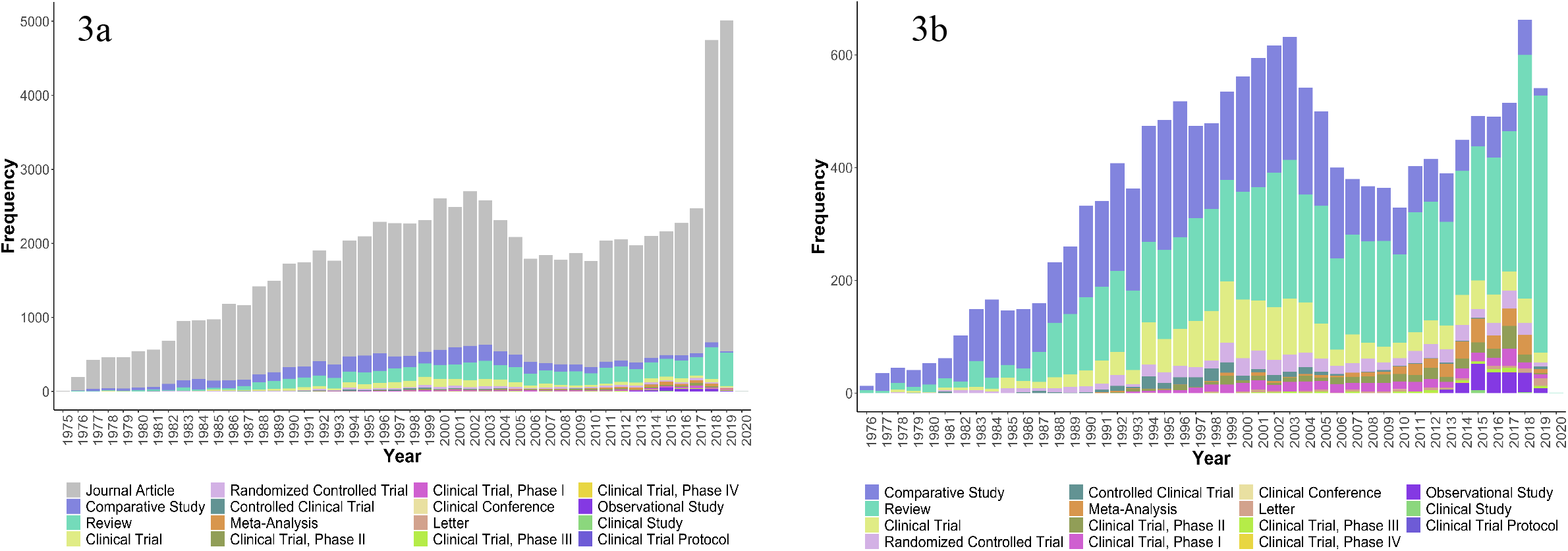
Nature and trends of HLA related publications in PubMed annually from 1975 onwards: Stacked Bar plot shows distribution of PubMed articles in different categories. a) Diverse studies including clinical trials are reported, with maximum numbers represented in the “journal article” category. b) A subplot of (a) after removing the most frequent “Journal article” type to visualise the trends in other categories.

### HLA genes, alleles and its distribution

There are 28,320 alleles with many associated with a disease or pathological conditions. There also exists a great extent of variability in the names within articles. E.g. HLA-B*13:01, a risk factor for dapsone hypersensitivity syndrome in multiple populations was written as HLA-B*13:01, HLA-B*1301, B*1301, B(*)1301 and B1301 in different papers. If one has to search for an allele and its related information, the user must be aware of all possible formats of writing an allele encompassing its current and previous nomenclature. So, based on this, we converted all existing HLA keywords to a standard allele name. We identified only ∼1% of the total alleles to be associated with conditions like diseases, graft survival, or drug reactions. To represent these alleles in the form of a graph, we collapsed the nomenclature to two-digit level (**Figure 4**). Majority of the studies were with HLA-DRB1 loci, followed by HLA-B and HLA-A, while fewer studies were on HLA-C locus. Each HLA alleles, collapsed to its two-digit information are linked to AFND server in the database, highlighting its allele frequency. The focus of our present study was also to understand the semantics between alleles and diseases, wherein we noted that some alleles were reported as protective and some as risk alleles. e.g. reports indicated HLA-DRB1*15 was protective for HIV and risk allele for pulmonary tuberculosis (28, 29). We were also interested in exploring the effects of multiple alleles individually on a single disease. To address this, we listed out 42 articles (**Supplementary Table 2**)highlighting the fact that for a single disease, different alleles can have contrasting effects, e.g. HLA-DQA1*02:01 and HLA-DQB1*06:02 can be protective in Artemisia pollen-induced allergic rhinitis while HLA-DQA1*03:02 can be a risk factor (30).

**Figure 4.**
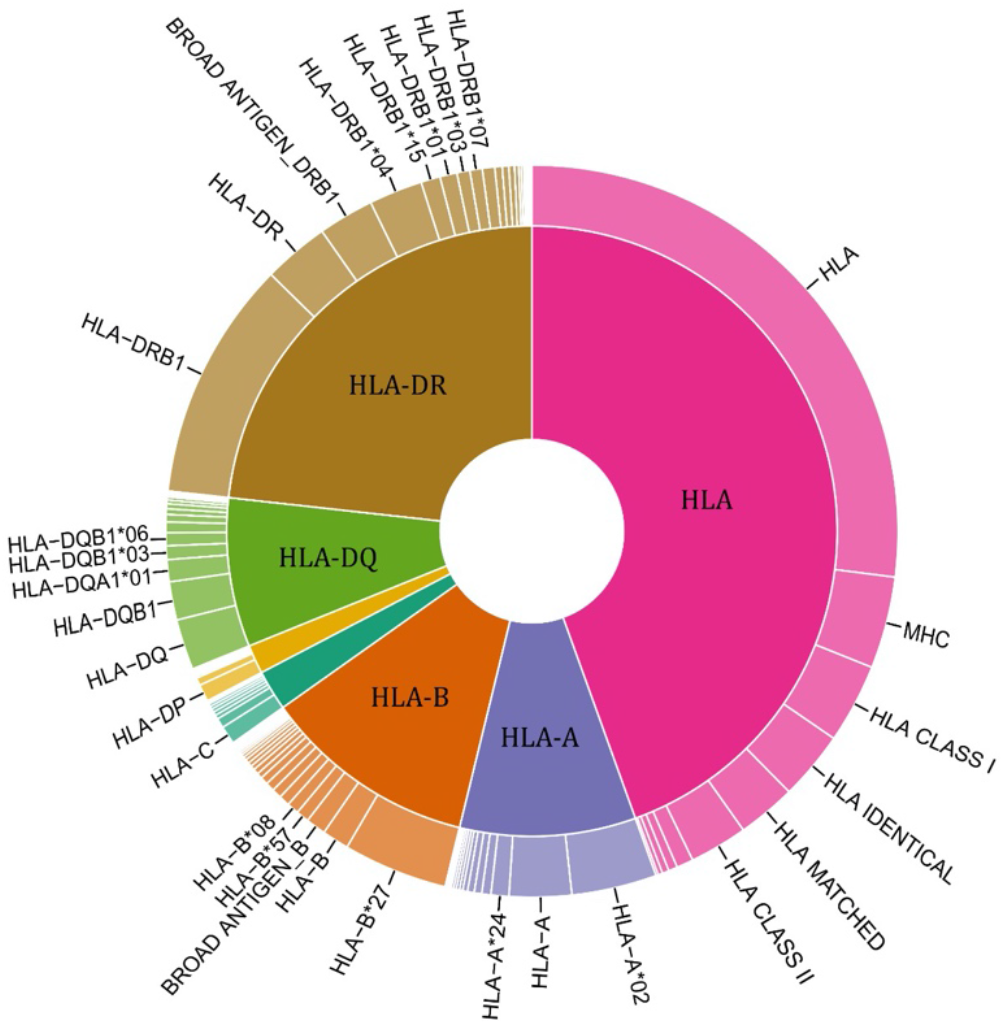
The topmost reported HLA alleles associated with diseases: All the HLA alleles indicated have been grouped to their second digit and represented in the pie chart. HLA-A, HLA-B and HLA-DRB1 are the most studied amongst the HLA genes.

### Exploring diseases, its associated categories and other relevant information

The HLA studies were divided into four broad categories: Diseases, Transplantations, Sign and Symptoms, and Therapeutics/ADRs, to study the information systematically. This grouping was done based on the MeSH keywords identified in the abstracts There are a total of 24 categories for diseases in MeSH, ranging from C01 and C04 through C26. We grouped C23 as “Sign and Symptoms” and C20.452 (GVHD) as part of Transplantation and rest as disease categories. Keywords falling under E04 (Transplantation procedures) were also grouped under “Transplantation”. For “Therapeutics/ADRs”, we selected only those sentences that had mentions of drug keywords, allele name and disease names together. We filtered them further if they satisfied either of the three conditions: 1) Belongs to Drug adverse reactions category or 2) Sentences had mentions of keywords such as reactions, -induced(carbamazepine-induced) or 3) Disease keyword had mention of –induced (Drug-induced liver injury). The remaining were grouped as “Diseases”. **Table 1** shows the number of articles under each category.

**Table 1:** Number of articles in broad categories

| **Categories** | **Number of PubMed abstracts** |
| --- | --- |
| Diseases | 29713 |
| Transplantation | 9258 |
| Signs and Symptoms | 6050 |
| ADR | 317 |

To study the association with diseases, we analysed data from both the “Diseases” and “Transplantation” category. Inconsistency in writing disease names increases the efforts in searching a specific query. To reduce this variability, MeSH ID was used to summarise the obtained information e.g. diseases like tumour, cancer, malignancy, and neoplasm (malignant and benign) were mapped to a single entity malignancy (D009369). Collapsing a large number of similar keywords to a single ID reduces the complexity in searching for articles related to particular diseases. We observed a total of 3615 different disease terms mapping to unique 1869 MeSH IDs. **Figure 5** represents a snapshot of common HLA associated diseases. To examine the disease associations, we mapped it to level-one (level-zero) terms. Diabetes Mellitus Type 1, Rheumatoid Arthritis, Multiple Sclerosis (Autoimmune Disease), Melanoma and Leukemic (Neoplasms by Histologic Type), Psoriasis (Skin disease) and Celiac Disease (Metabolic) were the topmost HLA associated diseases. In the analysed abstracts, the list of HLA associated diseases/conditions indicates that some diseases were very frequently reported, whereas other diseases like Down syndrome, Guillain-Barre Syndrome, Polymyalgia Rheumatica were infrequently or rarely reported. **Supplementary Table 3** represent the distribution of both common and less explored HLA associated diseases.

**Figure 5.**
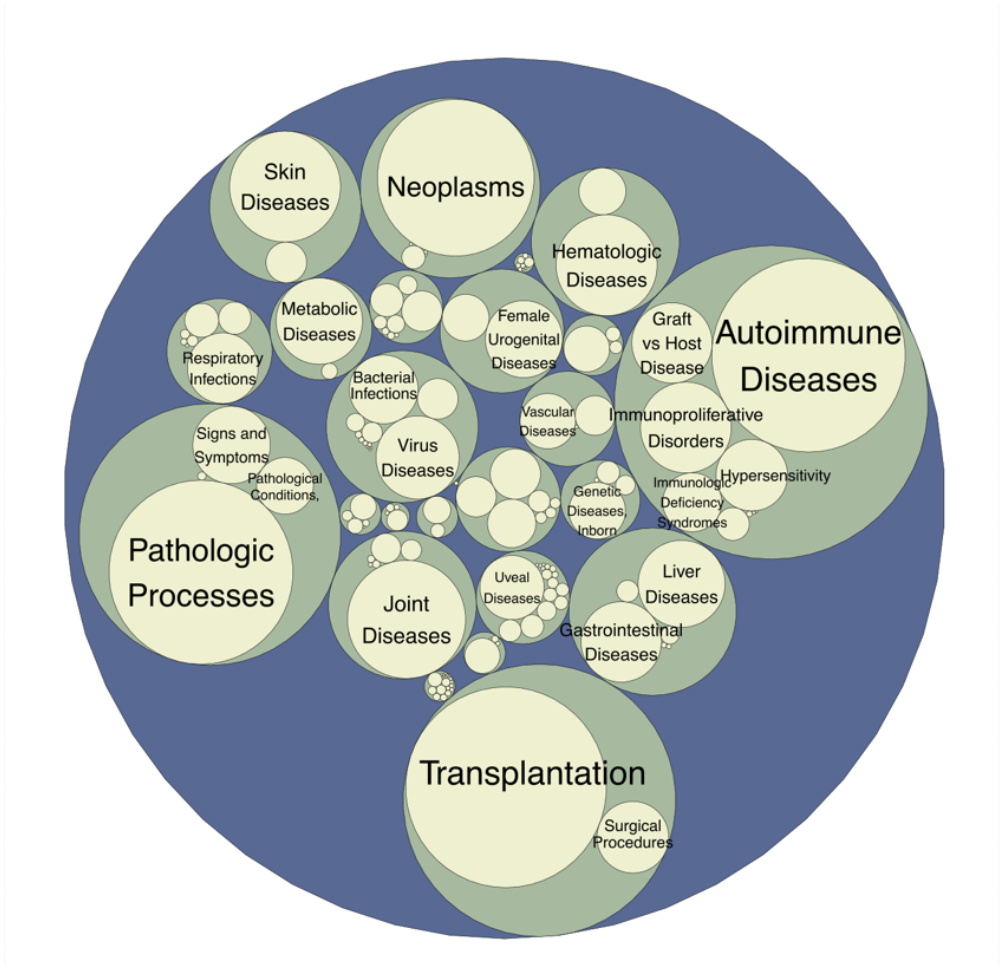
Diseases/conditions associated with HLA genes: Graph represents three level hierarchy of diseases. Each colour represents a level. There are 24 major categories as represented in green colour, which is further divided into subcategories. Each disease name is matched to its Mesh id and a normalised mesh keyword. Autoimmune, Neoplasms and Joint disease are the top most associated diseases. As anticipated, significant numbers of studies related to transplantation are also observed.

To get an overall perspective of genes and diseases, we considered the diseases at level-one along with HLA gene. We observed the majority of reported associations with HLA-DRB1, followed by HLA-B and HLA-A (**Figure 6**). We also listed details of individual allele-disease pairs for more information (**Supplementary Table 4**). HLA-DRB1 was reported to be linked with disease conditions like rheumatoid arthritis, type 1 diabetes, multiple sclerosis, melanoma and 1156 other diseases. HLA-B association was reported with spondylitis, infections, hypersensitivities, psoriasis, drug allergies and 859 other diseases and HLA-A was reported to be associated with melanoma, leukemia, influenza, haemochromatosis, and 705 other diseases.

**Figure 6.**
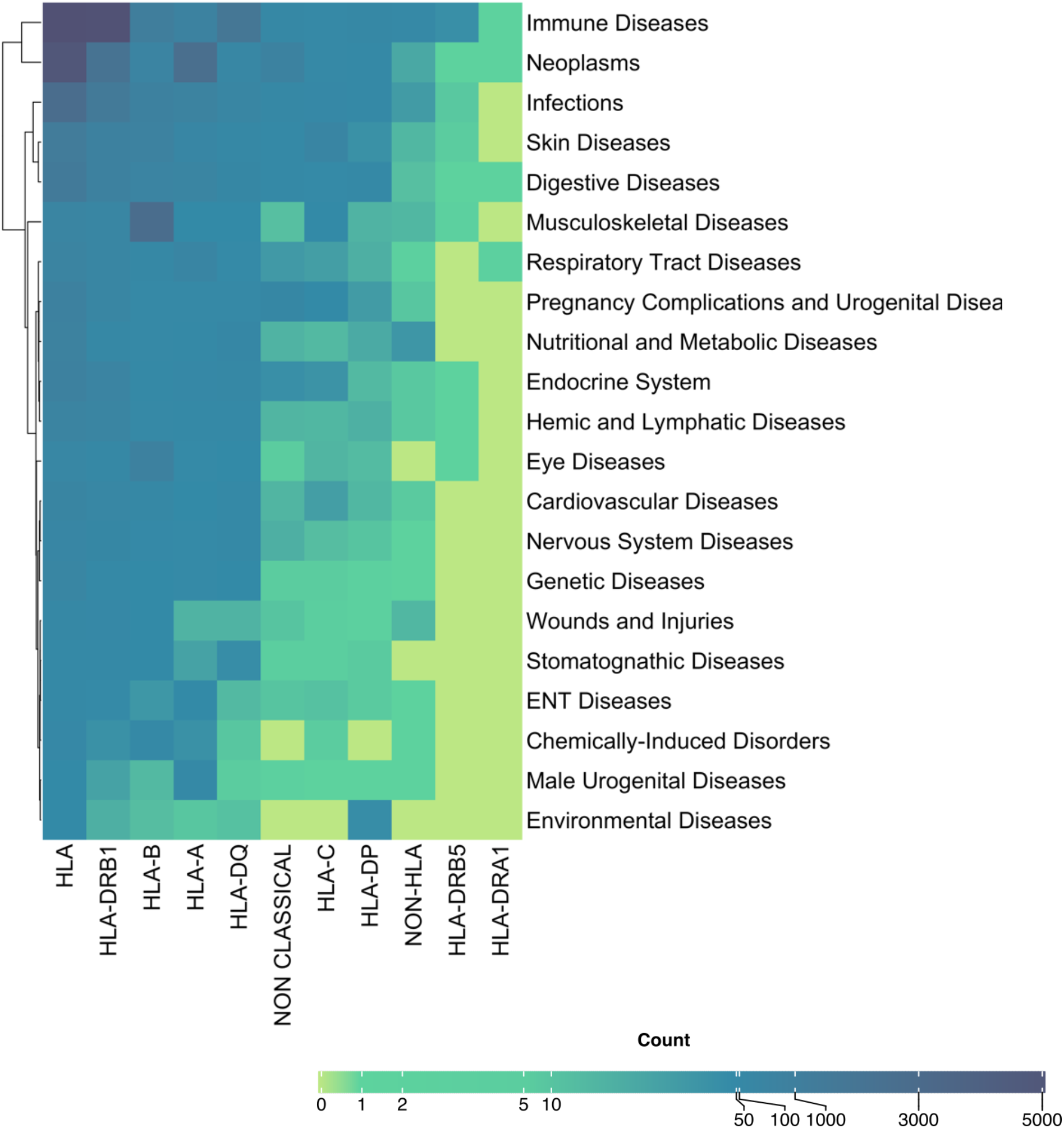
Heatmap of HLA Disease associations: The gradient heat map representing the number of diseases associated with HLA genes. First column represents generic “HLA” studies where specific gene information is not mentioned. A large number of associations were also observed with Non-classical(HLA-E,F,G) genes.

The analysis also takes into consideration the diseases which require transplantation and also include the complications associated with it both pre and post-transplantation. As anticipated, we observed that individuals suffering from beta Thalassemia and sickle cell anaemia (genetic and congenital disorders), multiple myeloma (an immunoproliferative disorder) and liver injury underwent transplantations of bone marrow, hematopoietic stem cells and renal tissue. However, there were other additional details included with the transplantation data such as disease history of patients before undergoing transplantation e.g. psoriasis, Graves’ disease, diabetic neuropathy and post-transplantation complications e.g. Ischemia, Necrosis, Fibrosis, Haemorrhage.” Such collated information under one platform may be of interest to a clinician for designing therapy modules. **Supplementary Table 5** represents details of transplantation related studies.

### SNPs and HLA diseases

HLA loci have a repertoire of genetic variations, a large number of which have been linked to multiple diseases via genome-wide association studies (GWAS). Though GWAS lists information about SNPs in/associated with HLA gene, a number of genetic variation studies go unnoticed either because they are small cohort analysis or are not compiled in a single resource for systematic study. Thus, to include the overlooked studies and missing information, this analysis reports information from all kinds of studies and includes abstracts mainly from journal articles, review, metanalysis, letters, and clinical trials. To acquire robust data, we retained only those HLA variations, that are present in the sentences along with the disease and allele keywords. We identified 313 unique SNPs mention and its details is compiled in **Supplementary Table 6**. Majority of SNPs mapped to intronic variants followed by missense and intergenic. **Figure 7** represents genomic distribution of mapped SNPs. A substantial number of variations also mapped to genes other than HLA, indicating they may be in Linkage Disequilibrium (LD) or frequently occur in conditions like transplantation success or ADRs example. We observed top hits of SNPs mapping to infectious diseases like HIV and hepatitis, inflammatory conditions like psoriasis, complex diseases like asthma and diabetes and hypersensitivity largely attributed by drug ADRs. SNP association studies are also based on a proxy SNP, which can be in LD with the causal variant and the LD values vary from one population to another. To address this, we also added population information of the studies whenever available in the abstract. The most studied SNP rs9277535, associated with hepatitis B virus, has been studied across a large number of populations from Asian and central Asian countries like China, Japan, Asia, Turkey, Korea, and Indonesia.

**Figure 7.**
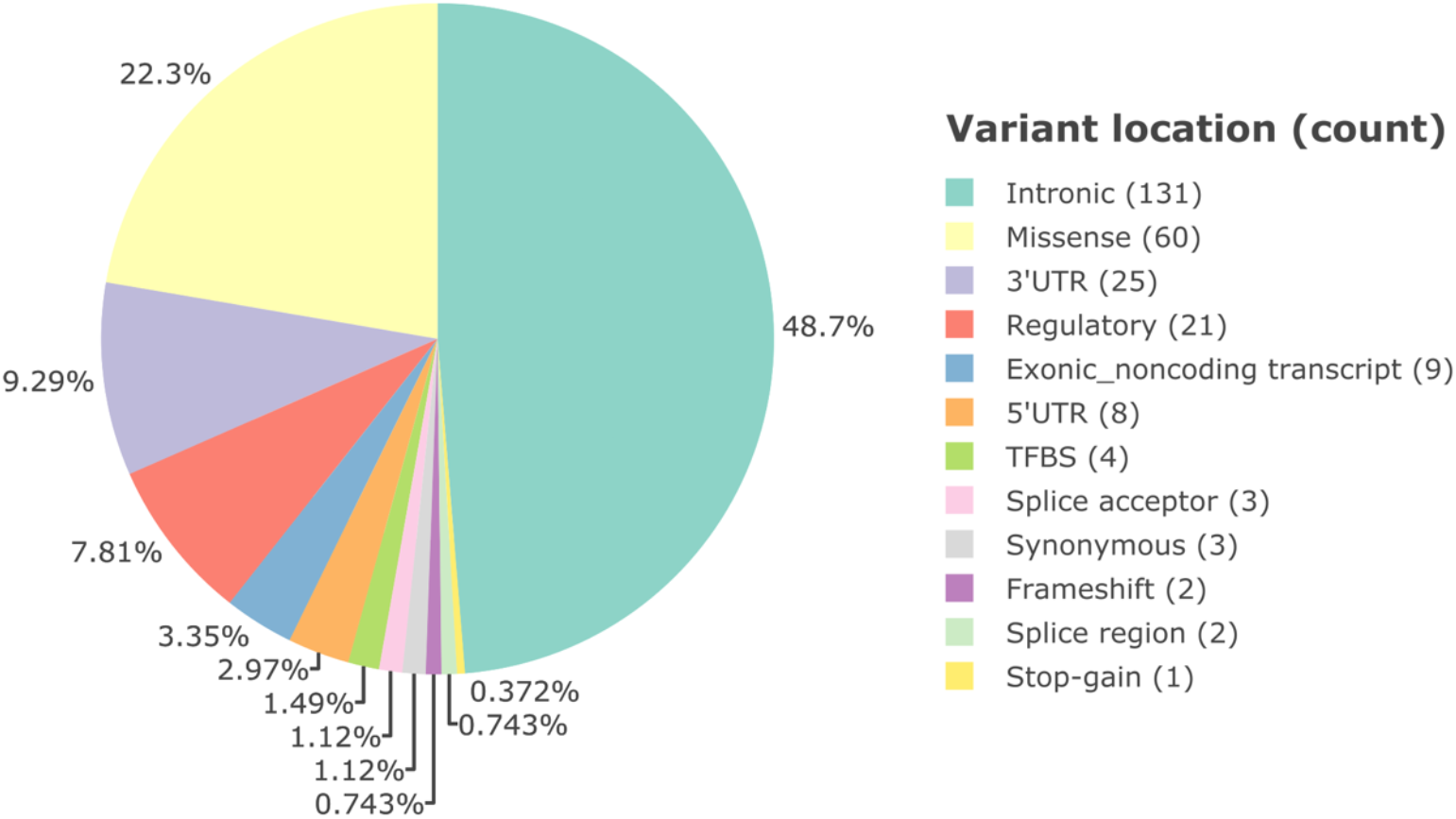
Genomic distribution of SNPs: Pie chart representing the number of variations in genic region with majority of them mapping to introns.

### Geographical Spread of HLA literature across various ethnic groups and populations

Genetic differences in HLA genes across populations and their link with biological conditions make it imperative to consider geographical information while studying HLA association with a particular condition. We assumed that the population/ethnic groups name might not be present in the same sentences that mention HLA and disease, so we used a flexible approach here and fetched the names of geographical locations present anywhere in the abstracts. In total, we reported 7696 NORP tags, mapping to 144 unique geographical entities. These unique tags were binned into 112 country-based populations and 32 ethnic groups. **Figure 8** represents the frequency distribution of these matched populations belonging to the countries and ethnic groups. Japan, China, USA, India and Italy are the major countries where the HLA gene-disease association studies have been reported with disease groups as shown in **Supplementary Table 7**. Along with this, the European subcontinent has been extensively studied (1129 reports) as a major ethnic group. Apart from frequently studied areas, we also observed locations like New Zealand, Armenia and Sri Lanka that have a low number of reported studies. This type of analysis can help researchers understand not only the extent of allele-disease associations among populations in the context of these immune players but also the scope of research in their selected geographical location while planning their hypothesis.

**Figure 8.**
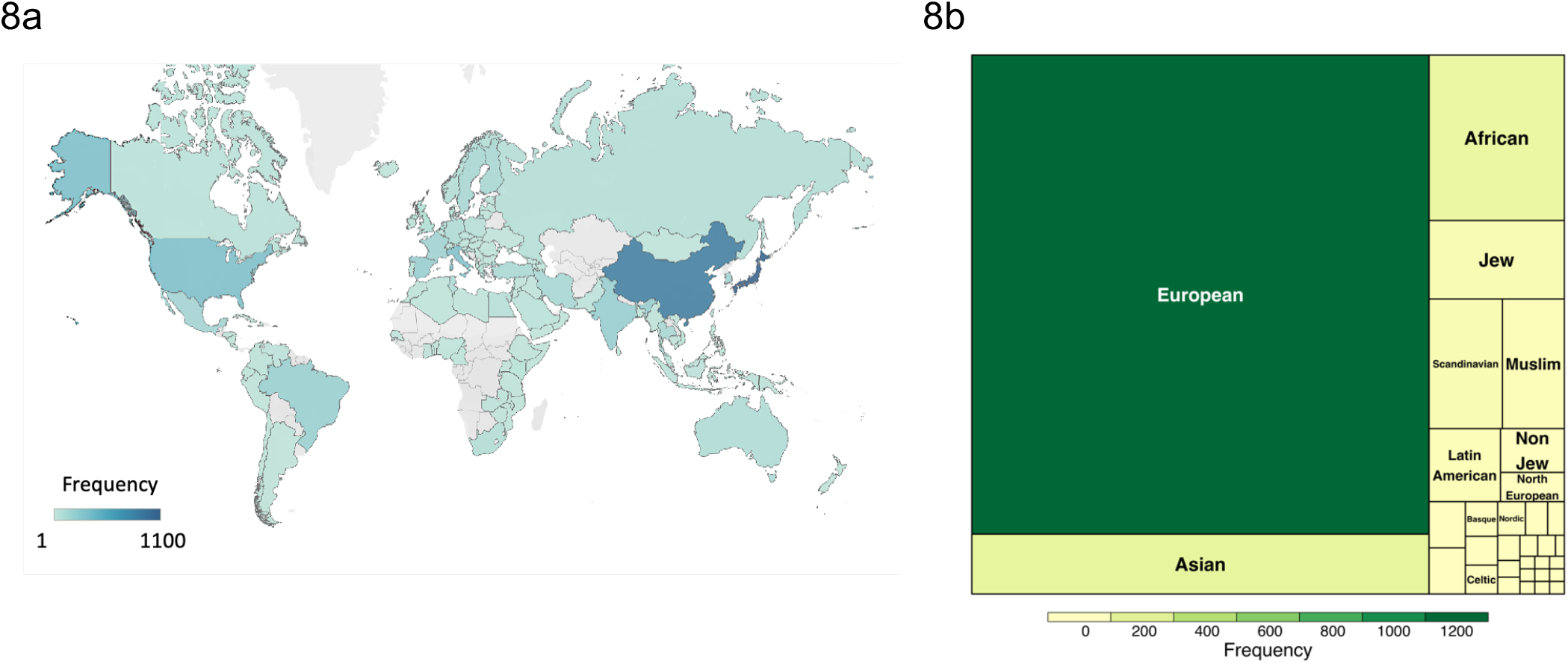
Geographical Spread of HLA studies: Identified geographical locations are binned to the nearest a) Country b) Ethnic group. Color gradient representing the count of various HLA alleles with respect to disease or ARD’s studies. China, Japan and the USA report maximum studies and European, Asian and African are the most studied ethnic groups

### Response to therapeutics

HLA genes are known to have association with various hypersensitivities and drug reactions, a few of them like Stevens-Johnson syndrome can also be life-threatening. Due to allele differences among individual and population level, these hypersensitivities vary, and thus studying these pharmacogenetic markers with the population information becomes important. For instance, we observed from our data that HLA-A*31:01 is associated with carbamazepine induced Stevens-Johnson syndrome in European population while HLA-B*15:02 is associated with Chinese and Indian populations. A meta resource like HLA-SPREAD can help understand such population-wise differences that obstruct designing of therapy modules for ADRs/ hypersensitivities. To be more specific, this analysis focuses on drugs that are present in sentences along with the disease and allele keywords. We observed a total of unique 317 abstracts mentioning 242 unique drugs, of which 52 mapped to ADR category. Details of drugs and related information are listed in **Supplementary Table 8**. We also validated our results with AFND, a manually curated database that has information about ADRs (**Figure 9a**). Out of 52 drugs present, we were able to find 33 common with AFND. One of the drugs “Valporic acid”, mentioned in AFND, was not present in the actual cited article and 8 drugs could not be captured because of the stringent criteria of drug mapping i.e. the drug name should be present in the sentence along with disease and allele keyword. **Figure 9b** lists the frequency-based distribution of top 20 drugs fetched from our analysis. Interestingly, we also observed 19 drugs that are not mentioned in AFND database, e.g. HLA-B*38:02:01 allele was found to predict carbimazole/methimazole induced agranulocytosis, HLA-DRB1 associated azathioprine induced pancreatitis in IBD patients. This analysis highlights, how one can miss information apart from the time and manpower intensive nature in manual curation.

**Figure 9.**
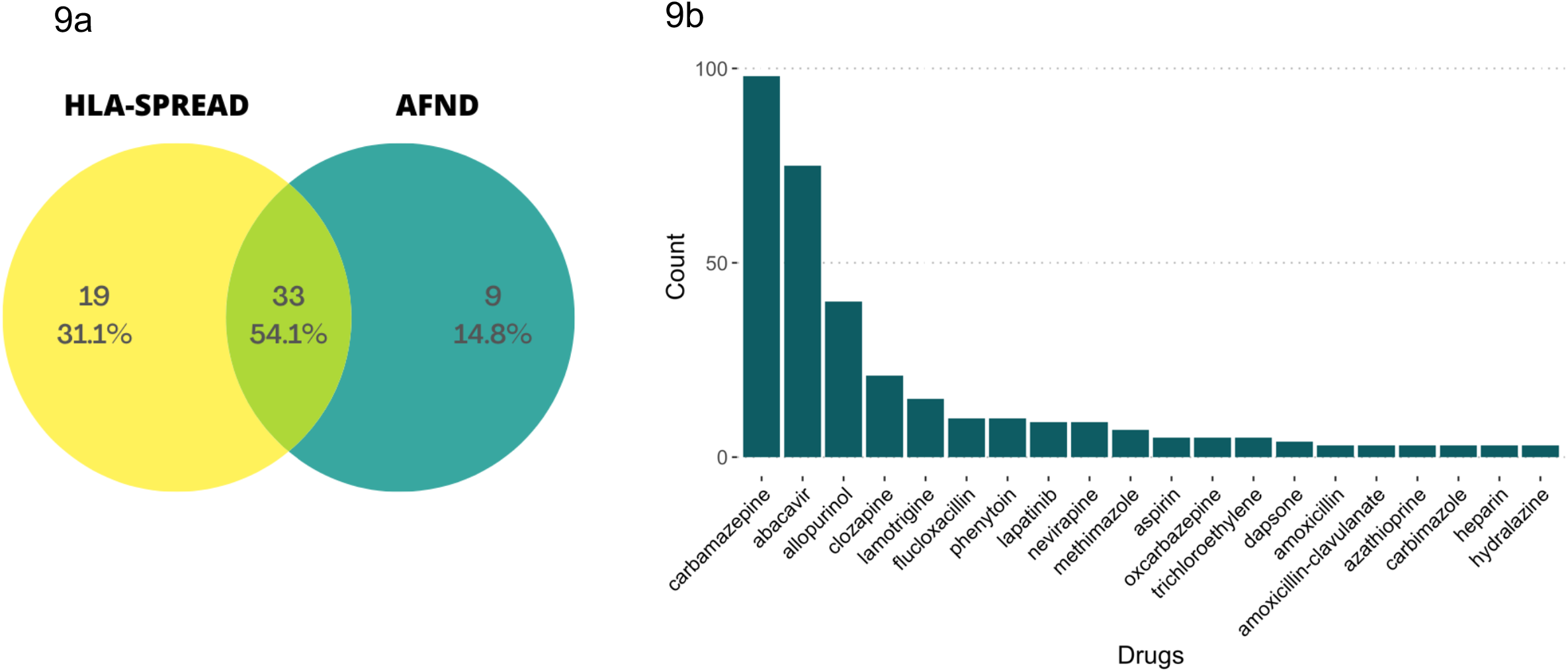
Statistics of drugs related HLA studies: A) Comparison of ADR’s identified using HLA-SPREAD with AFND. B) Bar plot showing the topmost 20 drugs identified.

### Insights from HLA-SPREAD: Biomarker ANALYSIS

We demonstrate the usability of the database to address clinically relevant queries. Multiple questions on the identification of HLA alleles and diseases linked with hypersensitivity, allergy, genetic marker, prognosis and diagnosis can be addressed using HLA-SPREAD. As an example, we present an analysis to identify biomarkers in HLA studies. To address this question, we used an n-gram based approach to identify the keyword most frequently occurring with “marker” in the sentences. **Table 2** list the most common keywords identified. We checked the details of such sentences and complied the information (**Supplementary Table 9**). A few of them like abacavir hypersensitivity and SJS syndrome were present in multiple papers. HLA-G and HLA-E were also reported to be markers for conditions like tumour, transplantation and heart diseases.

**Table 2:** Most frequently keyword occurring with “Marker”

|  |
| --- |
| Biomarker |
| Genetic marker |
| HLA marker |
| Predictive marker |
| Prognostic marker |
| Risk marker |
| Susceptibility marker |

## Discussion

HLA alleles are known to be associated with a large number of diseases. There is no existing repository that summarises this information in a systemic manner. Manual curation is a cumbersome process and one might also miss a lot of important information. The need for such a user-friendly platform increases significantly since HLA alleles have been found clinically associated with a large number of conditions. NLP based text mining offers a way to fetch this information pragmatically. NLP is instrumental in terms of extracting information from unstructured data. This method has started assuming immense importance in the biomedical domain. A few papers like GWASkb and SNP literature have used it for extracting information such as SNP and its related knowledge from the biomedical data whereas Monarch initiative has used it for studying phenotype information (31). Extracting information from HLA related literature is very difficult owing to the large number of studies and complex nomenclature. This project is an attempt to consolidate all the HLA relevant information such as SNPs, populations studied, ADRs and associated diseases into a structured database. This resource is also handy for user-specific advanced HLA searches like looking for biomarkers for toxicity-based studies and disease progression.

There were a few drawbacks of this analysis worth highlighting – primary arising due to the different formats of various journals. The initial tokenised data used in the analysis was based on English stop words. However, we observed in a small set of papers, the author missed giving full stops or spaces which lead to the fusion of two sentences. The subheadings were present in different cases and often followed by different special characters leading to complexity in their removal. Also, a prefix of keywords like SETTINGS, STUDY DESIGN, etc. have been observed in a few sentences, as those papers did not follow standard headlines. Apart from these, few other parameters like abbreviations at the end of sentences, presence of roman letters in sentences and different brackets and quotes styles in title caused errors during tokenisation process. Similarly, it was observed that with the updation of various abstracts in new releases, the previous incorrect entries were not removed which lead to duplication of different information.

Since HLAspread has catalogued information from diverse resources, in many instances it provides pieces of information that would be more informative and exhaustive. For instance, besides information retrieved from databases like DisGeNET, OMIM (Mendelian) reporting information on a few diseases we also used MESH is more comprehensive as it houses 139264 variant disease terms mapping to 4674 diseases. We also reduced the high variability in the method of mentioning the disease name in various articles. On average, a disease has around 30 names with one ID, showing the wide spectrum of disease dictionary required to capture all possible disease terms. In order to capture the HLA and ADRs we selected a list of drugs from SIDER4.1. However, not all drugs present in side effect database will be associated with ARDs. To get a more specific answer, we selected drugs from categories such as adverse drug reactions, hypersensitivity and toxicity. We were able to fetch a large number of studies and observed that the AFND database has missed quite some drugs in the ADR analysis. We thus added information from both AFND and SIDER to get heuristic information for a set of different drugs. There were a few unique aspects that we could capture because of our approach. For instance, in transplantation studies in addition to just listing different kinds of transplantations, we also observed the most common diseases which required transplantation and drugs given during the process with few side effects. Also, a unique aspect we added was a category called signs and symptoms for simplifying user searches. For instance, some users may also be interested in knowing the context of HLA alleles with conditions like inflammation, relapse, hypoxia, septic shock, diarrhoea, etc. We aim to add a few features in future updates for example mapping the variants reported in dbSNP, OMIM, ClinVar with to the HLA alleles. This would help in seamless integration of high-throughput variation data with the wealth of HLA information in literature and HLA alleles reported in IMGT database.

To summarise this is one of its kind of efforts to integrate the diversity of HLA information into a structured format for ease of query and analysis. This could also provide an informative resource for the non-HLA specialists for initiating any new studies in populations and diseases.

## Acknowledgements

The authors would Acknowledge COE M/o AYUSH grant MLP-901 to MM and DD and SRF fellowship to DD from Department of Biotechnology (DBT), and Dr. Yatender Kumar (NSIT) for permitting AK to work on this project. We would also acknowledge Mr Praveen Sinha for designing webpage of HLA SPREAD, Dr. Debasis Dash, CSIR-IGIB for critical reviewing of work, Dr. Ganesh Bagler and Rudransh Tunwani from IIITD for NLP discussion, Dr. Ganganath Jha from Hazaribagh University in QC of population curation and Malika Seth in QC of semantic annotations. The authors would also like to acknowledge Mr. Raghunandanan MV and Mr. Amit Khulve at CSIR-IGIB for IT support.

## Authors Contributions

MM, DD designed the study and co-wrote the manuscript. DD and AK executed the entire work. B.R.M developed the database. UK helped in HLA analysis, interpretation and manuscript writing.

## Supplementary tables

https://zenodo.org/record/4592338#.YEfBL10za3I

